# Oligodendrocytes depend on MCL-1 to prevent spontaneous apoptosis and white matter degeneration

**DOI:** 10.1101/2020.12.02.408138

**Authors:** Abigail H. Cleveland, Alejandra Romero-Morales, Laurent Alfonso Azcona, Melisa Herrero, Viktoriya D. Nikolova, Sheryl Moy, Orna Elroy-Stein, Vivian Gama, Timothy R. Gershon

## Abstract

Neurologic disorders often disproportionately affect specific brain regions, and different apoptotic mechanisms may contribute to white matter pathology in leukodystrophies or gray matter pathology in poliodystrophies. We previously showed that neural progenitors that generate cerebellar gray matter depend on the anti-apoptotic protein BCL-xL. Conditional deletion of *Bcl-xL* in these progenitors produces spontaneous apoptosis and cerebellar hypoplasia, while similar conditional deletion of *Mcl-1* produces no phenotype. Here, we show that, in contrast, postnatal oligodendrocytes depend on MCL-1. We found that brain-wide *Mcl-1* deletion caused apoptosis specifically in mature oligodendrocytes while sparing astrocytes and oligodendrocyte precursors, resulting in impaired myelination and progressive white matter degeneration. Disabling apoptosis through co-deletion of *Bax* or *Bak* rescued white matter degeneration, implicating the intrinsic apoptotic pathway in *Mcl-1*-dependence. *Bax* and *Bak* co-deletions rescued different aspects of the *Mcl-1*-deleted phenotype, demonstrating their discrete roles in white matter stability. MCL-1 protein abundance was reduced in *eif2b5*-mutant mouse model of the leukodystrophy vanishing white matter disease (VWMD), suggesting the potential for MCL-1 deficiency to contribute to clinical neurologic disease. Our data show that oligodendrocytes require MCL-1 to suppress apoptosis, implicate MCL-1 deficiency in white matter pathology, and suggest apoptosis inhibition as a leukodystrophy therapy.

## Introduction

Apoptosis is a highly conserved process across metazoans, with an increasing diversity of homologous regulatory proteins in more complex organisms. This diversity allows different types of cells to set different thresholds for apoptosis according to physiologic needs. In the brain, differentiated neurons, which are not replaceable through proliferation, down-regulate key apoptotic regulators such as *Apaf*, to become highly resistant to apoptosis (1, 2). In contrast, CNS progenitor cells, which are abundant and readily replaced through proliferation, rapidly undergo apoptosis in response to genotoxic stress. Genetic deletion studies show that specific brain progenitor populations are primed for apoptosis and depend on specific anti-apoptotic proteins to prevent spontaneous triggering of cell death mechanisms. Deletion of either of the anti-apoptotic homologs *Mcl-1* or *Bcl-xL (gene/transcript name BCL2L1)* causes spontaneous apoptosis in multi-potent precursor cells of the prenatal brain (3). In the postnatal brain, *Bcl-xL* dependence is associated with the neural lineage; deletion causes spontaneous apoptosis in differentiating cortical neural progenitors, resulting in microcephaly (4), and in cerebellar granule neuron progenitors, resulting in cerebellar hypoplasia (5). MCL-1 incompletely compensates for BCL-xL in cerebellar progenitors, as co-deletion of *Bcl2l1* and *Mcl-1* accelerates cerebellar growth failure, but isolated *Mcl-1* deletion in cerebellar granule neuron progenitors does not cause apoptosis or impair growth (6). Thus different types of brain cells, at different times and in different regions, show different mechanisms of apoptosis regulation, with specific developmental consequences.

Considering the importance of MCL-1 in the prenatal brain, and the BCL-xL-specific dependence of neural-committed cerebellar granule neuron progenitors in the post-natal brain, we investigated whether specific cells in the post-natal brain showed MCL-1 dependence. We deleted *Mcl-1* conditionally, using GFAP-Cre or NESTIN-Cre transgenic lines to target CNS stem cells in the prenatal brain that give rise to both neurons and glia. We found that the resulting *Mcl-1*-deleted mice showed normal brain anatomy and neurologic function through postnatal day 7 (P7) but then developed progressive postnatal white matter degeneration and increasing ataxia. The symptoms, white matter specificity, and early postnatal onset *of the Mcl-1* deletion phenotype resembled clinical leukodystrophies observed in patients. Genetic co-deletions that blocked the intrinsic apoptotic pathway rescued the *Mcl-1* deletion phenotype, confirming the role of apoptosis in the pathogenesis of MCL-1-deficient white matter degeneration, and suggesting that impaired apoptosis regulation through MCL-1 may contribute to clinical white matter disorders.

## Methods

### Mice

We generated *Mcl-1*^*cKO*^ mice with conditional, brain-specific *Mcl-1* deletion, by crossing *GFAP-Cre* mice (Jackson Labs, stock # 004600) that express Cre recombinase in neuro-glial stem cells of the brain with *Mcl-1*^*LoxP/LoxP*^ mice (7), generously shared by Dr. You-Wen He. We interbred GFAP-Cre, *Mcl-1*^*loxP/loxP*^, and *Bax*^*LoxP/LoxP*^*/Bak*^*-/-*^ (Jackson Labs, stock # 006329) to generate *Mcl-1;Bax*^*dKO*^, *Mcl-1;Bak*^*dKO*^, and various heterozygous controls. All mice were of species *Mus musculus* and crossed into the C57BL/6 background through at least five generations. All animal studies were carried out with the approval of the University of North Carolina Institutional Animal Care and Use Committee under protocols (19-099).

Wild type (C57BL/6 strain) and *eIF2B5*^*R132H/R132H*^ mice (8) were generated, bred, and housed in Tel Aviv University animal facility (Lab Products Inc., Seaford, DE, USA). All experimental procedures were approved by the Tel Aviv University Animal Care Committee according to national guidelines (permit #04-17-022). Breeding and genotyping were performed as previously described. (8) Each generation was established by back cross of homozygous *eIF2B5*^*R132H/R132H*^ mice with WT C57BL/6J (Harlan Labs, Jerusalem, Israel) to prevent genetic drift.

### Histology and immunohistochemistry (IHC)

Mouse brains were processed and immunostained as previously described (9-12). In brief, mice were decapitated under isoflurane anesthesia and brains were harvested and drop fixed in 4% formaldehyde for 24 hours, then transferred to graded ethanol series embedded in paraffin and sectioned. Eyes were processed and fixed as previously described (13). In brief, eyes were harvested after decapitation, injected with 4% formaldehyde, and then drop fixed, embedded in paraffin and sectioned.

Primary antibodies used were: BAK diluted 1:200 (Cell Signaling, #12105), MBP diluted 1:1000 (Abcam, #ab7349), SOX10 diluted 1:100 (Santa Cruz, #sc-17342), cleaved-Caspase 3 (cC3) diluted 1:400 (Biocare Medical, #CP229C), glial fibrillary acidic protein (GFAP) diluted 1:2000 (Dako, Z0334), PDGFRA diluted 1:200 (Cell Signaling, #3174), SOX2 1:200 (Cell Signaling, #4900S), SOX9 1:200 (R&D Systems, #AF3075), NESTIN 1:500 (Cell Signaling, #4760), IBA1 1:2000 (Wako Chemicals, #019-19741). Stained images were counterstained with DAPI, digitally imaged using an Aperio Scan Scope XT (Aperio) and subjected to automated cell counting using Tissue Studio (Definiens).

### Magnetic resonance imaging

For MRI studies, brains from P7, P14, and P21 mice were harvested after decapitation, fixed in 4% formaldehyde in PBS for 48hrs, then transferred to PBS. Brains were then embedded in 2% agarose in 15mL plastic vials and imaged in pairs as previously described (14).

### Western blot

Whole brains or cerebella were harvested and homogenized in lysis buffer (Cell Signaling, #9803). Lysate protein concentrations were quantified using a bicinchoninic acid (BCA) protein assay kit (Thermo scientific, #23225). Equal total protein concentrations were resolved on SDS-polyacrylamide gels (BioRad, # 4561105, # 4568094) and transferred onto polyvinylidene difluoride membranes. Membranes were blotted using a SNAP i.d. 2.0 Protein Detection System (Millipore). Membranes were imaged using a chemiluminescent SuperSignal West Femto Maximum Sensitivity Substrate (34095, Thermo Fisher Scientific) and the C-DiGit blot scanner (LI-COR Biosciences). Signal was quantified using the Image Studio Lite software (LI-COR). The following antibodies were used for Western blot: MCL-1 (Cell Signaling, #94296; 1:500 dilution), BAX (Cell Signaling, #14796; 1:500 dilution), β-actin (Cell Signaling, #3700; 1:5000 dilution), Actin (Sigma #A3853; 1:240000 dilution), Anti-rabbit IgG, HRP-linked antibody (Cell Signaling, #7074; 1.5:1000 dilution), Anti-mouse IgG, HRP-linked antibody (Cell Signaling, #7076; 1.5:1000 dilution). For the eIF2B5^R132H/R132H^ studies, cerebrums were harvested from 7 and 10 month old mice and protein was extracted using a lysis buffer containing 1% triton, 0.5% NaDOC, 0.1% SDS, 50 mM Tris pH 8, 100 mM NaCl, 10 mM β-glycerophosphate, 5 mM NaF, 1 mM DTT, 1 mM vanadate, and EDTA-free complete TM protease inhibitor cocktail (#11-836-170-001; ROCHE). Protein was quantified using a BCA protein assay kit (#23227 Pierce), diluted in 1X sample buffer (0.05 M HEPES pH 7.4, 2% (w/v) SDS, 10% (v/v) Glycerol, traces of Bromophenol Blue, 100 mM Dithiothreitol (DTT)), and separated on a 10% SDS-PAGE gel. Blots were imaged using an AI600 Imager (Amersham) and quantified by ImageQuant TL (GE Healthcare, Pittsburgh, PA, United States).

### Behavioral studies

#### Animals included

Subjects were *Mcl-1*^*cKO*^ mice or littermate controls with intact *Mcl-1*, including genotypes Mcl1^f/+^ without Cre, Mcl1^f/f^ without Cre, and Gfap-Cre/Mcl1^f/+^. A total of 8 litters were needed to generate all of the replicates. Day of birth was considered postnatal day 0 (P0). Subject numbers at the beginning of the study (P7) were 13 (21%) *Mcl-1*^*cKO*^ and 48 (79%) control. All litters completed the first 2 weeks of testing. Three litters (n=23) were removed from the study for histology assays during the third week of testing: one on P15, after the second open field test; one on P16, after the first acoustic startle test; and one on P21, after the tail suspension test.

#### Behavioral testing regimen

#### Developmentally-calibrated behavioral tests

For motor tests at P7, mice were removed from the home cage and maintained in a beaker warmed to 35°C. Mice were then subjected to the neurobehavioral negative geotaxis, righting, and open field tests, adapted from published studies (15).

#### Negative geotaxis

A 20 cm × 20 cm screen with 0.5 cm wire mesh squares was set at a 25° angle. Each pup was placed downward on the screen with its head facing toward the bottom, allowed to grip the mesh, and then released. The time it took for each pup to turn 180° (head and upper torso vertical), was recorded, with a 30 s maximum and each pup was tested 3 times.

#### Righting

Each pup was placed onto its back on a flat Plexiglas surface. The experimenter gently held the pup in place for 2 s and then released it. The latency to righting (all 4 feet on the ground) was timed, with a maximum of 30 s.

#### Open field activity and pivoting

Each pup was placed into the center of a Plexiglas chamber (PhenoTyper; 30.5 cm × 30.5 cm × 43.5 cm; Noldus Information Technology, Wageningen, The Netherlands). A dim yellow light (30 lx) at the top of the PhenoTyper box was illuminated throughout the 3 min trial. Measures were taken of distance traveled and time in the center region (4 cm × 4 cm square in the center of the arena) by an image-tracking system (EthoVision, Noldus Information Technology). In addition, the experimenter recorded the number of vertical rearing movements and number of pivots (circling or lateral movement, as opposed to forward locomotion).

Each mouse was screened through the 3 assays in the order presented here. After finishing the neonatal motor screen, each pup was weighed, marked with a non-toxic Sharpie, and placed back into the warm beaker. The litter was returned to the home cage after all pups had been tested.

#### Open field activity at P14 and P19

Additional open field activity tests were performed at P14 and P19. The distance traveled, the number of vertical rearing movements, and time spent in the center region was recorded in each 10 min test.

#### Acoustic startle responses

This test was used to assess auditory function, reactivity to environmental stimuli, and sensorimotor gating. Mice were evaluated at 2 time points, P14 and P21. The procedure was based on the reflexive whole-body flinch, or startle response, that follows exposure to the acoustic stimuli. Measures were taken of startle magnitude and prepulse inhibition, which occurs when a weak stimulus leads to a reduced startle in response to a subsequent louder noise.

Mice were placed into individual Plexiglas cylinders within larger, sound-attenuating chambers. Each cylinder was seated upon a piezoelectric transducer, which allowed vibrations to be quantified and displayed on a computer (San Diego Instruments SR-Lab system). The chambers included a ceiling light, fan, and a loudspeaker for the acoustic stimuli. Background sound levels (70 dB) and calibration of the acoustic stimuli were confirmed with a digital sound level meter (San Diego Instruments). Each session began with a 5-min habituation period, followed by 42 trials of 7 different types: no-stimulus (NoS) trials, trials with the acoustic startle stimulus (AS; 40 msec, 120 dB) alone, and trials in which a prepulse stimulus (20 msec; either 74, 78, 82, 86, or 90 dB) occurred 100 ms before the onset of the startle stimulus. Measures were taken of the startle amplitude for each trial across a 65-msec sampling window, and an overall analysis was performed for each subject’s data for levels of prepulse inhibition at each prepulse sound level (calculated as 100 - [(response amplitude for prepulse stimulus and startle stimulus together / response amplitude for startle stimulus alone) × 100].

#### Limb clasping

At P18-21, a subset of mice (n=39) was assessed for limb clasping during a tail-suspension test. Each mouse was lifted by the base of the tail, and suspended above cage bedding for 15 sec, with ventral (stomach) side facing the experimenter. Response to tail suspension was coded using the following scoring system:

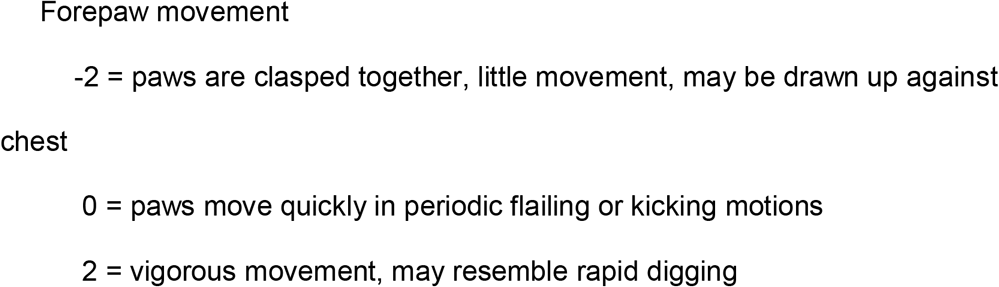

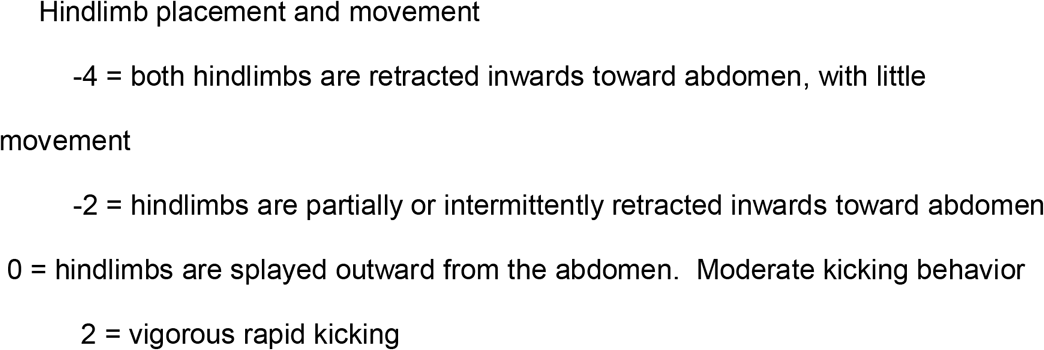

### Statistical analysis of behavioral tests

All testing was conducted by experimenters blinded to mouse genotype. Statview (SAS, Cary, NC) was used for data analyses. To reduce the number of *Mcl-1*^*cKO*^ subjects required, behavioral data from males and females were combined. One-way or repeated measures analysis of variance (ANOVA) were used to determine effects of genotype. Post-hoc analyses were conducted using Fisher’s Protected Least Significant Difference (PLSD) tests following a significant ANOVA F value. For all comparisons, significance was set at p<0.05.

## Results

### Conditional *Mcl-1* deletion in the CNS causes progressive white matter degeneration

We generated mice with conditional, brain-specific *Mcl-1* deletion by breeding *Mcl-1*^*loxP/loxP*^ mice that harbor loxP sites around exon 1 of the *Mcl1* locus (7), with *GFAP-Cre* mice that express Cre recombinase in stem cells that give rise to the neurons and glia of the cerebrum and cerebellum, excluding the Purkinje cells (Supplementary fig. 1A-B) (16, 17). The resulting *hGFAP-Cre/Mcl-1*^*loxP/loxP*^ (*Mcl-1*^*cKO*^) mice were born at the expected Mendelian ratio. We compared *Mcl-1*^*cKO*^ mice to littermate controls, including *Mcl-1*^*loxP/loxP*^ mice without Cre and heterozygously deleted *hGFAP-Cre/Mcl-1*^*loxP/+*^. We found no differences between control genotypes and therefore pooled them for analysis. In contrast, *Mcl-1*^*cKO*^ mice appeared normal at P7 but developed increasing ataxia, could not be weaned, and died between P20-23.

Examination of the brains of *Mcl-1*^*cKO*^ mice showed progressive white matter degeneration, that began after the first week of life, with relative sparing of gray matter. At P7, *Mcl-1*^*cKO*^ brains showed no abnormalities and appeared similar to controls (Fig. 1A, P7). At P14, *Mcl-1*^*cKO*^ mice showed white matter rarefaction and ventricular dilation and these changes were more pronounced at P21 (Fig. 1A, P14&21). In addition to white matter loss, we also noted migration abnormalities in the cerebellum, with ectopic cerebellar granule cells in the molecular layer of the cerebellum (Fig. 1A, P14&21). Replacing *Gfap-Cre* with *Nestin-Cre*, which similarly drives recombination in multipotent glio-neuronal stem cells, generated mice with the genotype *Nestin-Cre/Mcl-1*^*loxP/loxP*^ which displayed similar white matter-specific degeneration (Supplementary Fig. 1C). Deletion of *Mcl-1* throughout the brain thus resulted in progressive white matter pathology that began in the early postnatal period and increased over time.

**Figure 1.**
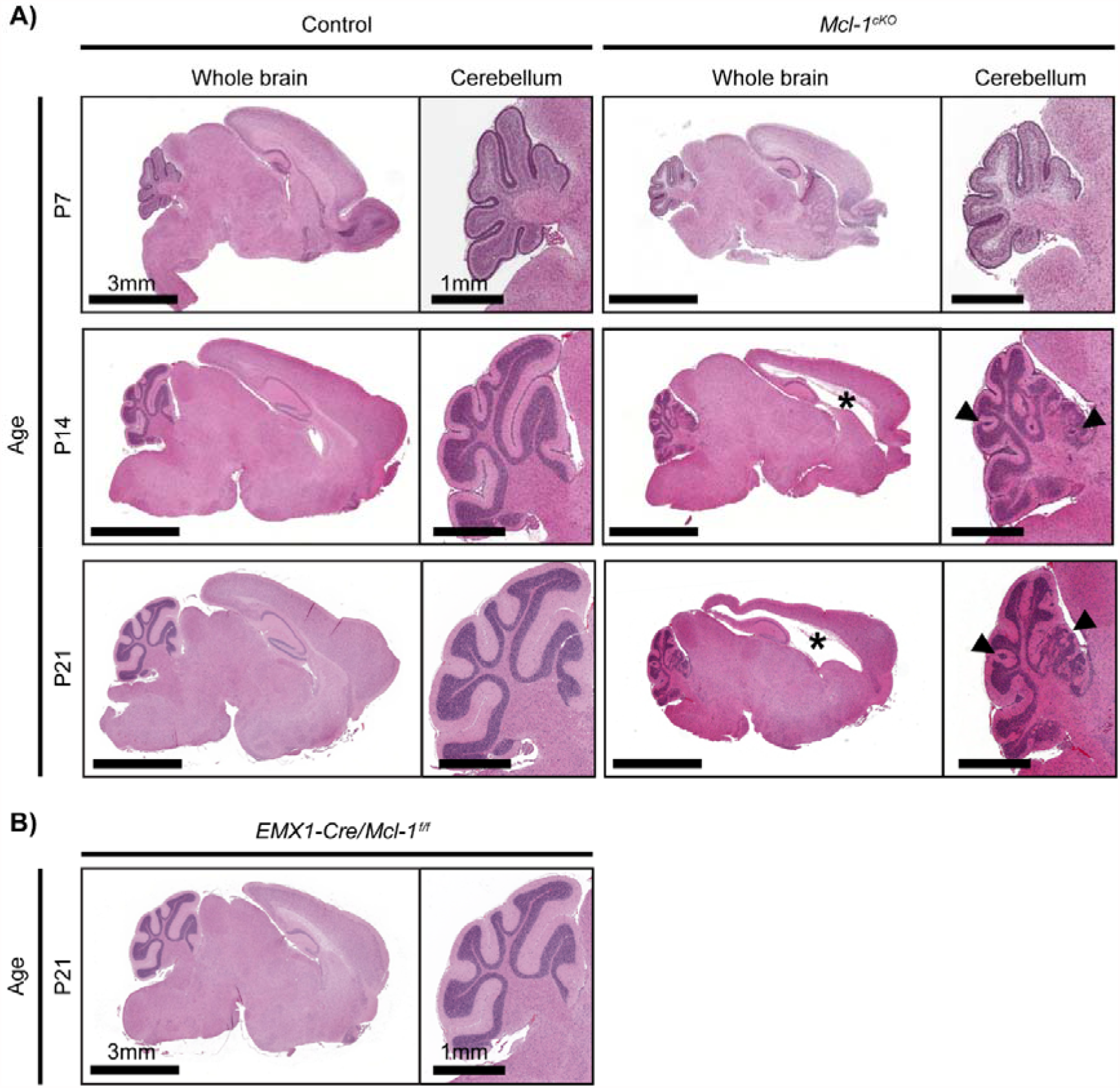
*Mcl-1* deletion causes progressive white matter degeneration. Brains of representative *Mcl-1*^*cKO*^ mice shown in H&E-stained sagittal sections appear similar to controls at P7. By P14, *Mcl-1*^*cKO*^ mice develop white matter degeneration with ventricular enlargement (*) and abnormal migration of cerebellar granule neurons (arrows). White matter loss increases by P21.

Consistent with the onset of neuropathology after P7, *Mcl-1*^*cKO*^ mice showed no behavioral deficits during the first week of life, then showed failure to thrive in the second week of life (Fig. 2A). Concurrent with reduced weight gain, *Mcl-1*^*cKO*^ mice demonstrated progressive abnormalities of motor function. We used developmentally adjusted tests to compare behavior of *Mcl-1*^*cKO*^ mice and controls at P7, P14, and P18. To examine behavior at P7, when mice do not yet have open eyes and do not venture from the nest, we used a test of geotaxis by placing mice on a tilted screen and measuring latency to turn. We also measured latency to turn over in a righting test, and we counted pivots during a 3-min open field test. We found no genotype-specific differences at P7 (Table 1). As P7 pups are beginning to gain control of hind limb function, open field testing allowed us to compare the number of pivots and distance traveled, which were similar in *Mcl-1*^*cKO*^ and control mice (Fig. 2B).

**Figure 2.**
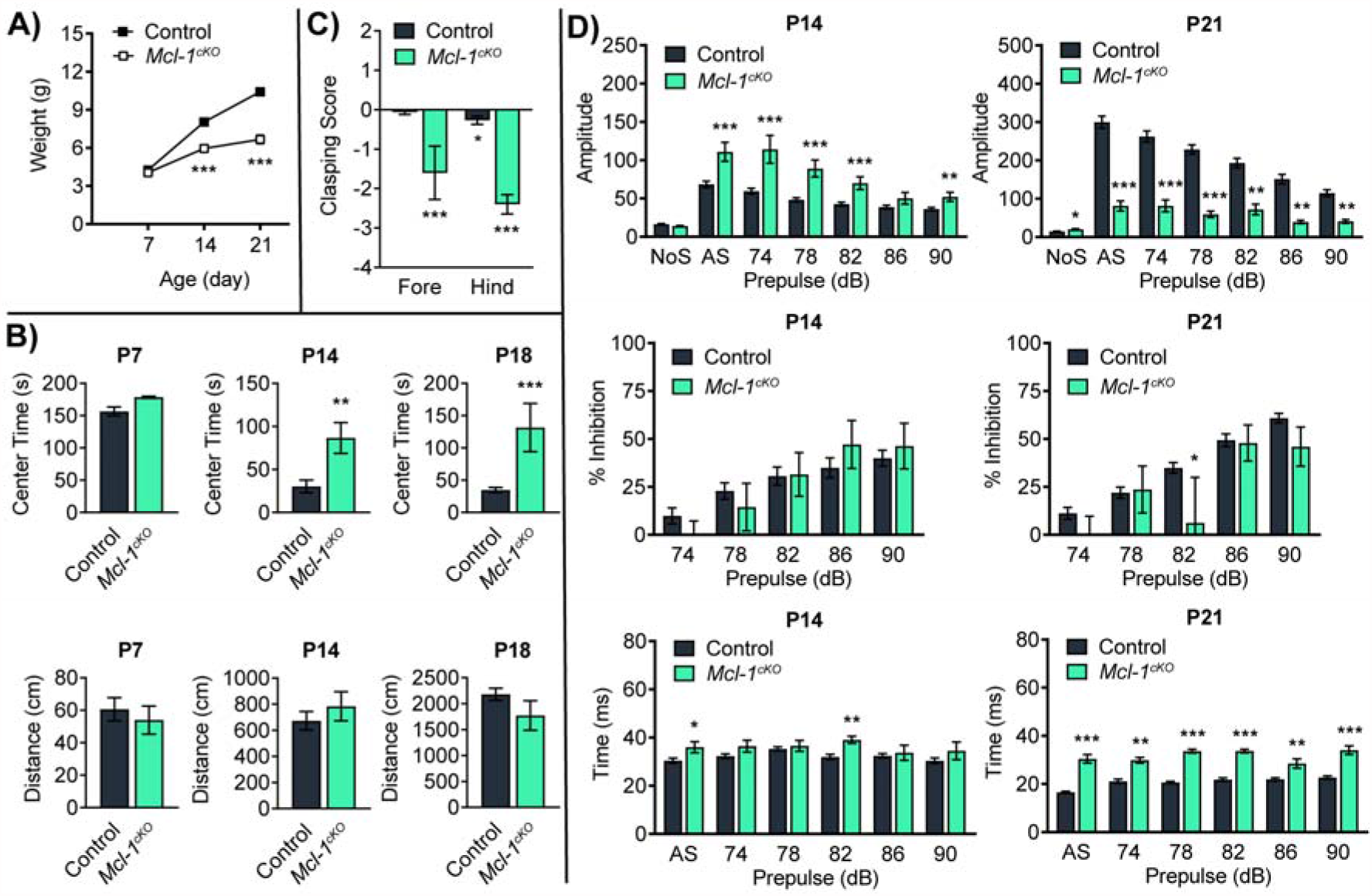
*Mcl-1* deletion causes progressive neurologic impairment. (**A**) Reduced weight gain in *Mcl-1*^*cKO*^ mice compared to controls. (**B**) *Mcl-1*^*cKO*^ mice show normal motor function at P7, with altered motor function by P14 and (**C**) increased hind-limb clasping at P18. (**D**) *Mcl-1*^*cKO*^ mice show increased startle in response to auditory stimuli at P14, and reduced elicited movement at P21. *, ** and *** denote p<0.05, p<0.01 and p<0.001 respectively, relative to controls.

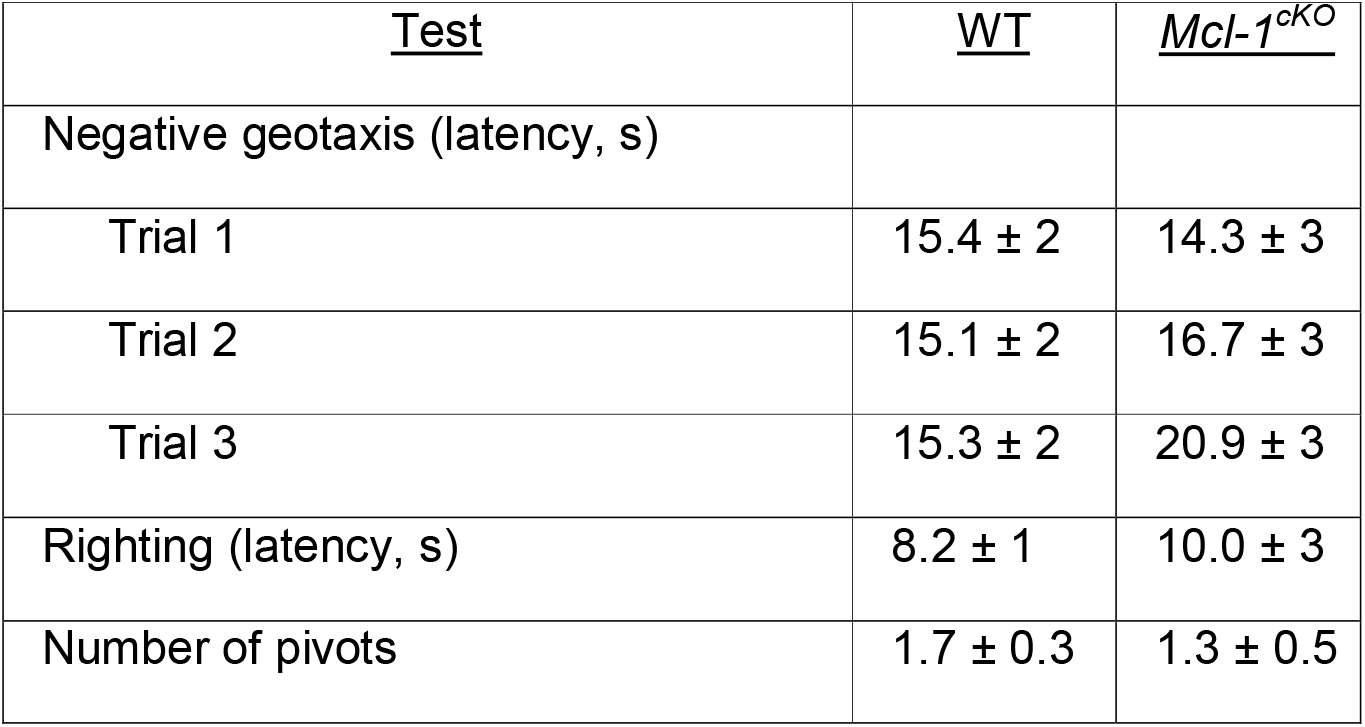

| <u>Test</u> | <u>WT</u> | <u>*Mcl-1<sup>ckO</sup>*</u> |
| --- | --- | --- |
| Negative geotaxis (latency, s) |  |  |
| Trial 1 | 15.4 ± 2 | 14.3 ± 3 |
| Trial 2 | 15.1 ± 2 | 16.7 ± 3 |
| Trial 3 | 15.3 ± 2 | 20.9 ± 3 |
| Righting (latency, s) | 8.2 ± 1 | 10.0 ± 3 |
| Number of pivots | 1.7 ± 0.3 | 1.3 ± 0.5 |

In contrast to the similar behaviors at P7, by P14 *Mcl-1*^*cKO*^ mice showed decreased rearing when placed in an open field and spent more time in the center of the open field, while traveling similar distances compared to controls (Fig. 2B; Supplementary Fig. 2A). Considering the differences in open field avoidance at P14, we examined whether *Mcl-1* deletion in *Mcl-1*^*cKO*^ mice produced retinal pathology that might confound behavioral analysis. We found no detectable pathology in the retinas or optic nerves of P15 *Mcl-1*^*cKO*^ mice (Supplementary Fig. 2B), and loss of vision was thus unlikely to explain reduced open field avoidance in *Mcl-1*^*cKO*^ mice. By P18, *Mcl-1*^*cKO*^ mice showed a marked increase in limb clasping when vertically suspended (Fig. 2C), a commonly used marker of disease progression in a number of mouse models of neurodegeneration and ataxia (4, 18).

Responses to sensory input were also abnormal in *Mcl-1*^*cKO*^ mice and changed over time. At P14, mice showed increased amplitude of acoustic startle response (Fig. 2D); these exaggerated responses demonstrated intact hearing but abnormal processing. By P21, when motor function was significantly impaired, the amplitude of acoustic startle response was markedly diminished compared to controls, and it is possible that impaired hearing may have contributed to this late abnormality. These motor and sensory tests showed progressive behavior abnormalities that correspond temporally with the white matter degeneration, beginning at P12 and worsening by P18-21.

### Blocking apoptosis prevents white matter degeneration in *Mcl-1*-deleted mice

To determine if white matter degeneration in *Mcl-1*^*cKO*^ mice resulted from increased apoptosis, we examined cell death through cleaved caspase-3 immunohistochemistry and TUNEL assays. We did not detect an increase in cleaved caspase-3-stained cells at P7, prior to the onset of white matter loss (data not shown), or at P11, when white matter loss was on-going (Supplementary Fig. 3, middle column). However, TUNEL+ nuclei of apoptotic cells are cleared more slowly than cC3+ apoptotic bodies and TUNEL may therefore be more sensitive for detecting increased cell death. TUNEL staining at P11 demonstrated increased cell death in *Mcl-1*^*cKO*^ brains, consistent with a pro-apoptotic effect of *Mcl-1* deletion (Fig. 3A).

**Figure 3.**
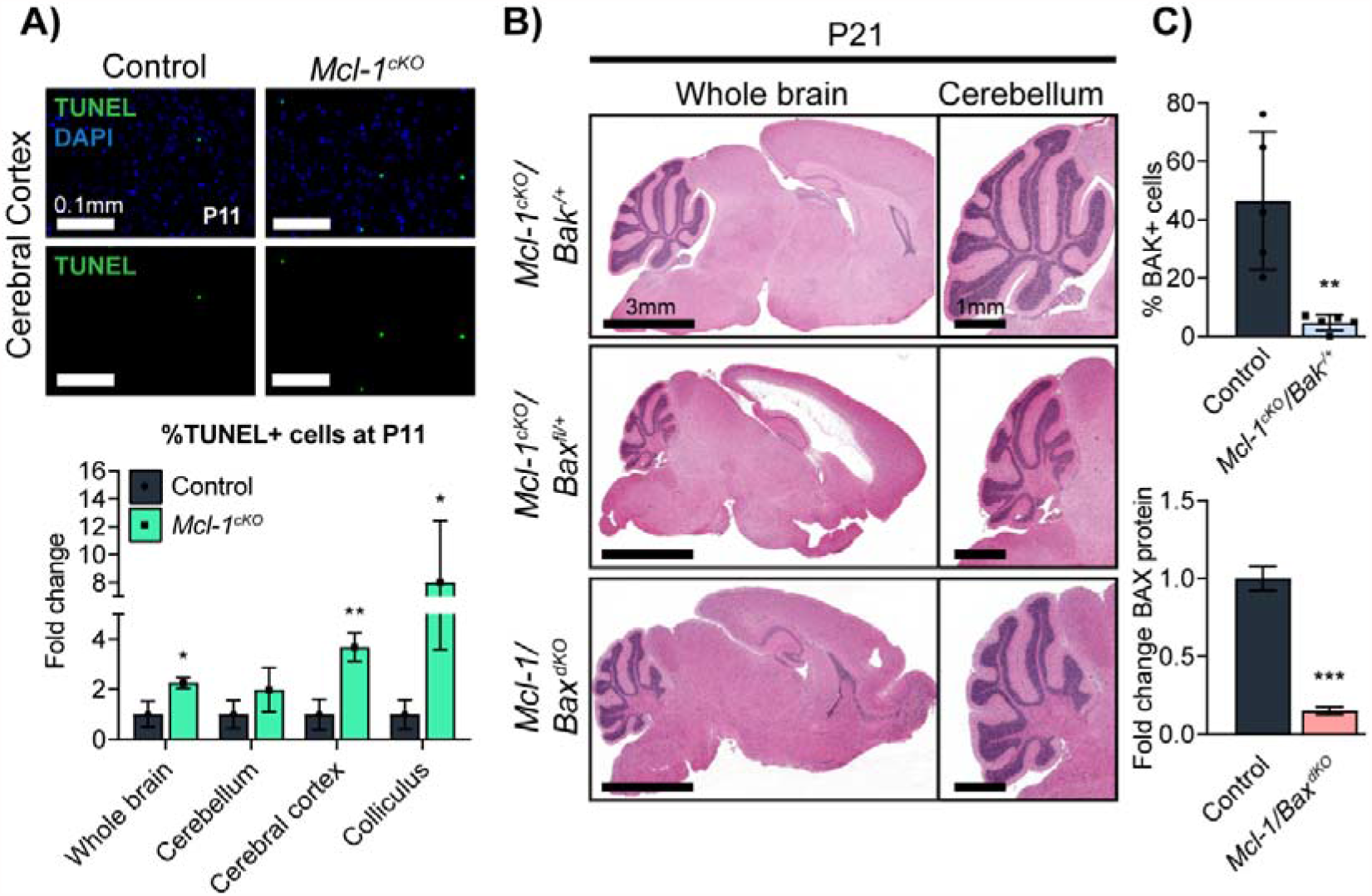
*Mcl-1* deletion causes white matter degeneration by increasing apoptosis. (**A**) TUNEL demonstrates increased cell death in *Mcl-1*^*cKO*^ brains. For cell counts, one sagittal section from each replicate was scanned at 20x and TUNEL+ cells were counted across the entire portion of each designated region that was contained within the section. (**B**) Representative H&E sections show that heterozygous *Bak* co-deletion and homozygous *Bax* co-deletion both rescue the white matter loss with ventricular enlargement (*) and cerebellar abnormalities caused by *Mcl-1* deletion. (**C**) Quantification of BAK immunohistochemistry in corpus callosum shows that heterozygous *Bak* deletion significantly reduces the number of BAK+ cells. Quantification of western blot of whole cerebella shows that homozygous *Bax* deletion reduces BAX protein abundance. *, ** and *** denote p<0.05, p<0.01 and p<0.001 respectively, relative to controls.

To determine whether the increased cell death in *Mcl-1*-deleted mice was causally related to the white matter degeneration, we tested whether blocking apoptosis through co-deletion of either *Bax* or *Bak* rescued the *Mcl-1*^*cKO*^ phenotype. We repeatedly intercrossed *Mcl-1*^*cKO*^ mice with *Bax*^*loxP/LoxP*^*/Bak*^*-/-*^ mice to generate animals with conditional deletion of *Mcl-1* and heterozygous or homozygous deletion of *Bax* or *Bak*. We then raised pups of each genotype and analyzed survival, neurologic function, and neuropathology.

We found that co-deletion of *Bax* or *Bak* significantly suppressed the *Mcl-1*^*cKO*^ phenotype, but resulted in different, mitigated phenotypes. *Mcl-1*-deleted mice with heterozygous co-deletion of *Bak*, with the genotype *hGFAP-Cre/Mcl1*^*loxP/loxP*^*/Bak*^*-/+*^ (*Mcl-1*^*cKO*^*/Bak*^*-/+*^), showed normal survival without overt neurologic deficits and normal white matter and cerebellar migration (Fig. 3B). In contrast, *Mcl-1*-deleted mice with heterozygous co-deletion of *Bax*, with the genotype *hGFAP-Cre/Mcl1*^*loxP/loxP*^*/Bax*^*loxP/+*^ (*Mcl-1*^*cKO*^*/Bax*^*fl/+*^), showed a progressive leukoencephalopathy phenotype similar *Mcl-1*^*cKO*^ mice (Fig. 3B). Immunohistochemistry demonstrated markedly lower BAK protein in *Bak*^*+/-*^ animals (Fig. 3C), indicating that that heterozygous *Bak* deletion was sufficient to alter BAK expression. The rescued phenotype of *Mcl-1*^*cKO*^*/Bak*^*-/+*^ mice therefore demonstrated reducing BAK protein expression was sufficient to prevent white matter degeneration.

*Mcl-1*-deleted mice with homozygous *Bax* co-deletion, with the genotype *hGFAP-Cre/Mcl1*^*loxP/loxP*^*/Bax*^*loxP/loxP*^ (*Mcl-1/Bax*^*dKO*^) showed normal survival, similar to *Mcl-1*^*cKO*^*/Bak*^*-/+*^ mice. The extensive white matter loss in *Mcl-1*^*cKO*^ mice was rescued in *Mcl-1/Bax*^*dKO*^ mice, as in *Mcl-1*^*cKO*^*/Bak*^*-/+*^ mice (Fig. 3B). The rescued phenotypes of *Mcl-1*^*cKO*^*/Bak*^*-/+*^ and *Mcl-1/Bax*^*dKO*^ mice show that disrupting either *Bax* or *Bak* is sufficient to prevent the severe white matter degeneration that otherwise results from *Mcl-1* deletion, implicating apoptosis in the pathogenesis of the *Mcl-1*^*cKO*^ phenotype.

### Mcl-1 deletion phenocopies human leukodystrophies

Leukoencephalopathies in patients produce characteristic MRI findings, including high water signal in white matter tracts (19), and the replacement of white matter by cerebrospinal fluid (20). We analyzed whether *Mcl-1*^*cKO*^ mice showed similar white matter changes on MRI. Similar to patients with the leukodystrophy VWMD, *Mcl-1*^*cKO*^ mice showed progressive ventriculomegaly (Fig. 4A). The increase in ventricle size was not evenly distributed, however, as the volume of the 4^th^ ventricle was significantly smaller in *Mcl-1*^*cKO*^ mice at P21 (Fig. 4B). Increased lateral ventricles and reduced 4^th^ ventricle was consistent with loss of periventricular cerebral white matter and an overall reduction in intracranial pressure, indicating hydrocephalus *ex vacuo*.

**Figure 4.**
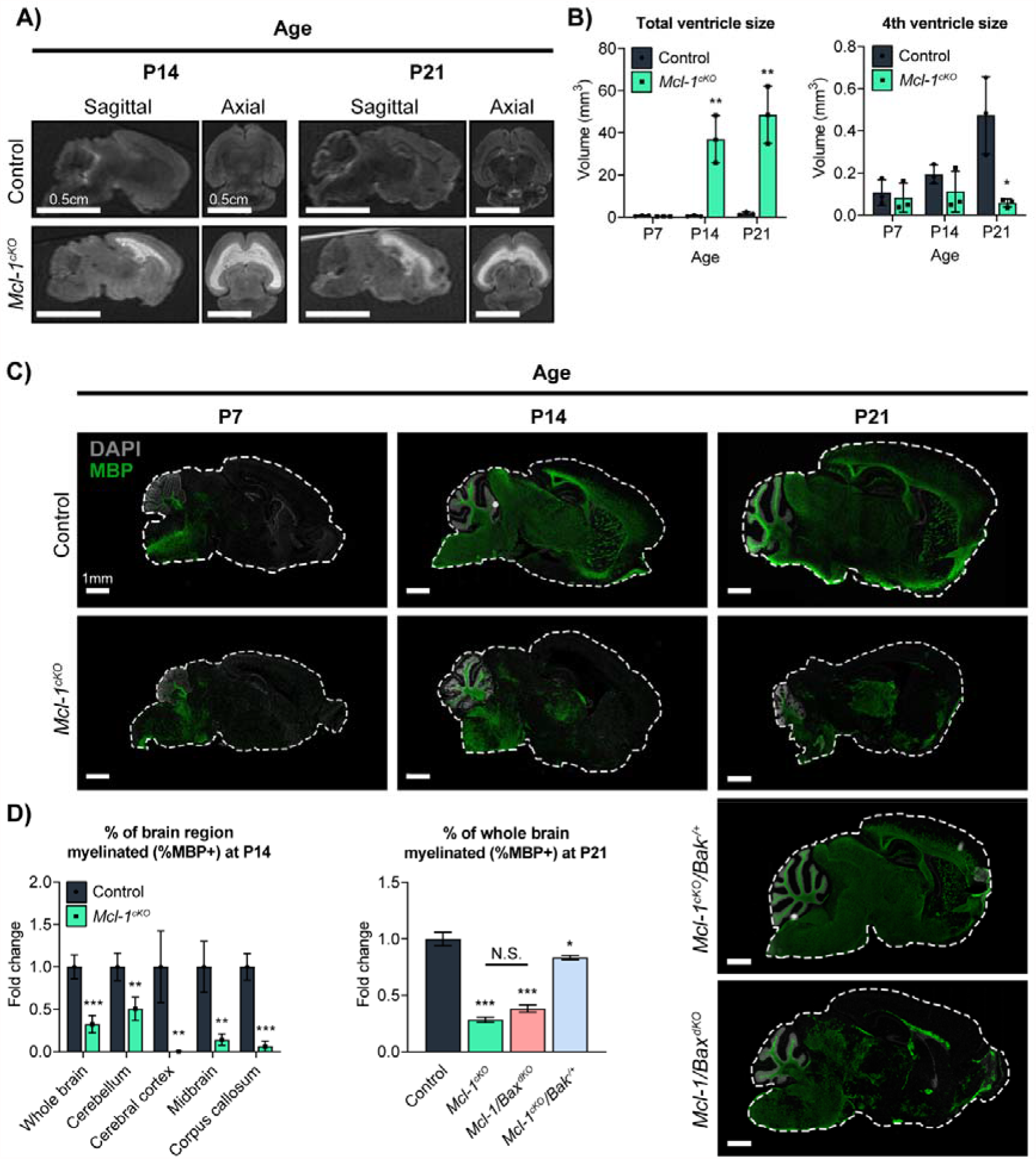
Abnormal neuroimaging and myelination in *Mcl-1*-deleted mice, with different rescue effects caused by co-deletion of *Bak* or *Bax*. (**A**) Representative MRIs show white matter hyperintensity in *Mcl-1*^*cKO*^ mice. (**B**) Total ventricular volume is abnormal in *Mcl-1*^*cKO*^ mice and increases from P7-P21, while the volume of the 4^th^ ventricle decreases significantly by P21. (**C**,**D**) Representative images and quantitative analysis of MBP IHC show that myelination is markedly decreased in *Mcl-1*^*cKO*^ mice, rescued in *Mcl-1*^*cKO*^*/Bak*^*+/-*^ mice, and incompletely rescued in *Mcl-1/Bax*^*dKO*^ mice. **(D)** Graphs show MBP+ area/total area within individual sections, normalized to the mean value in the control mice. *, ** and *** denote p<0.05, p<0.01 and p<0.001 respectively, relative to controls.

To determine whether myelination was disrupted in *Mcl-1*^*cKO*^ mice, we analyzed the distribution of Myelin Basic Protein (MBP). In control brains, the MBP+ area increased markedly from P7 to P14 (Fig. 4C). In contrast, myelination was delayed in *Mcl-1*^*cKO*^ brains, which showed significantly less MBP+ area at P14 in all brain regions analyzed (Fig. 4D).

Analysis of MBP+ area in *Mcl-1*-deleted mice with *Bax* or *Bak* co-deletions showed that a specific role for BAK in regulating survival of myelinating cells. We compared brains in P21 animals, when normal brains show widespread myelination. Heterozygous *Bak* co-deletion largely rescued myelination in *Mcl-1*^*cKO*^*/Bak*^*+/-*^ mice. In these animals, MBP+ area was over 2-fold greater than in *Mcl-1*^*cKO*^ mice, although we noted a small, statistically significant decrease in myelination compared to *Mcl-1* intact controls. Homozygous *Bax* deletion, however, did not rescue myelination in *Mcl-1/Bax*^*dKO*^ mice (Fig. 4D). The differential effects of *Bak* or *Bax* deletion on myelination support the interpretation that BAX and BAK act in cells with different functions, with both BAX and BAK regulating cells required for white matter maintenance, and only BAK regulating cells required for myelination.

### *Mcl-1* deletion reduces oligodendrocyte and Bergmann glia populations

To determine if impaired myelination in *Mcl-1*^*cKO*^ mice resulted from oligodendrocyte loss, we compared the numbers of cells expressing the oligodendrocyte markers SOX10 and PDGFRA in *Mcl-1*^*cKO*^ and control mice (Fig. 5A). SOX10 marks the oligodendrocyte lineage from the initial fate commitment in oligodendrocyte precursors (OPCs) to the maturation of myelinating oligodendrocytes, while PDGFRA specifically marks OPCs (21, 22). SOX10 cell counts thus reflect the total oligodendrocytic population across the differentiation spectrum, and PDGFRA cell counts reflect the undifferentiated subset.

**Figure 5.**
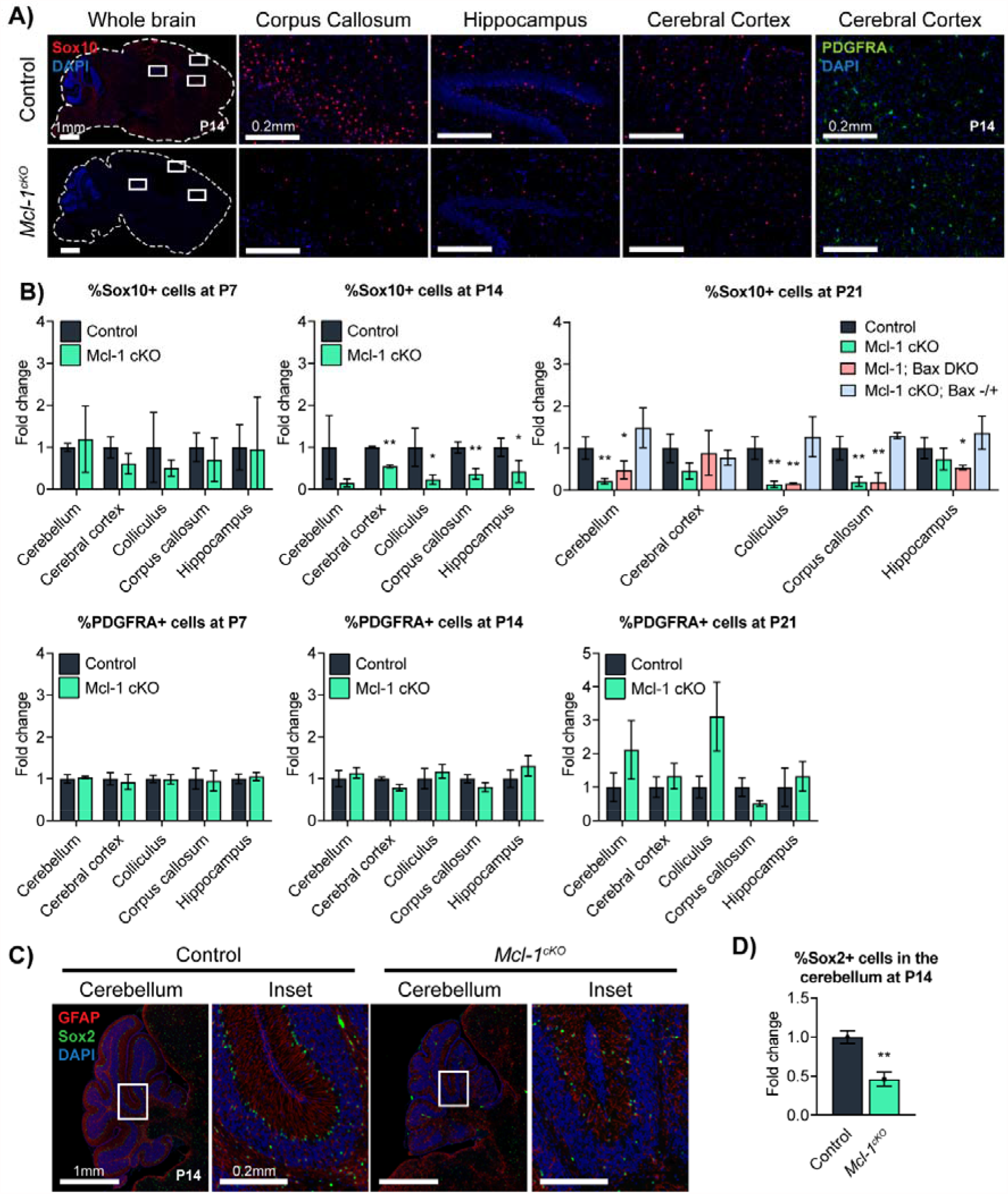
*Mcl-1* deletion causes loss of oligodendrocytes and Bergmann glia that is rescued with regional variation by co-deletion of *Bak* or *Bax*. (**A**,**B**) Representative images and quantification of SOX10 and PDGFRA IHC in the indicated genotypes. (**C**) Representative immunofluorescence for GFAP and SOX2 IHC identify Bergmann glia in P14 mice of indicated genotypes. (**D**) SOX2+ cells are significantly reduced in *Mcl-1*^*cKO*^ mice. * and ** denote p<0.05 and p<0.01 respectively, relative to controls.

At P7, oligodendrocyte populations in *Mcl-1*^*cKO*^ mice were not significantly different from controls. By P14, *Mcl-1*^*cKO*^ mice showed significantly fewer SOX10+ cells in diverse brain regions, while PDGFRA+ cells were similar between genotypes (Fig. 5B). These data indicating a global decrease in differentiated oligodendrocytes with preservation of OPCs. At P21, oligodendrocytic populations remained reduced in the cerebellum, corpus callosum, and superior colliculus; the lack of statistically significant differences in the cerebral cortex and hippocampus likely reflect the confounding effects of severe volume loss in these regions. In contrast to reduced SOX10+ cells, PDGFRA+ cells showed no decreases and rather trended toward increased populations at P21. Together our studies of SOX10 and PDGFRA show that oligodendrocyte loss in *Mcl-1*^*cKO*^ mice occurred specifically with differentiation.

To determine if apoptosis mediates the decline in the oligodendrocyte populations in *Mcl-1*-deleted mice, we examined the effect of *Bax* or *Bak* co-deletion in *Mcl-1*^*cKO*^*/Bak*^*+/-*^and *Mcl-1/Bax*^*dKO*^ mice. Immunofluorescence using antibodies to cC3 and SOX10 at P11 demonstrated a trend of increase in apoptosis in oligodendrocytes (p=0.06; Supplementary Fig. 3). Heterozygous *Bak* co-deletion in *Mcl-1*^*cKO*^*/Bak*^*+/-*^ mice normalized oligodendrocyte population size (Fig. 5B). In contrast, oligodendrocyte populations decreased in *Mcl-1/Bax*^*cKO*^ mice with homozygous *Bax* deletion (Fig. 5B). *Bak* co-deletion thus rescued the effects of *Mcl-1* deletion on white matter volume, myelination, and total oligodendrocyte populations. However, *Bax* co-deletion rescued only white matter volume, but myelination and oligodendrocyte numbers remained affected in *Mcl-1/Bax*^*dKO*^ mice. The discordant effects of *Bak* and *Bax* co-deletions indicates that different types of white matter cells depend on MCL-1 to control BAK-dependent or BAX-dependent apoptosis.

In contrast to the discordant effects of *Bax* and *Bak* co-deletions in oligodendrocytes, Bergmann glia were rescued by either co-deletion. We examined Bergmann glia because of the migration abnormalities in *Mcl-1*^*cKO*^ mice, as Bergmann glia guide cerebellar granule neuron progenitors as they migrate to the internal granule cell layer to become cerebellar granule neurons (23, 24). We identified Bergmann glia as cells with SOX2+ nuclei and GFAP+ processes (Fig. 5C). At P7, the numbers of Bergmann glia in *Mcl-1*^*cKO*^ were similar to control mice (data not shown), but by P14, the Bergmann glia populations in *Mcl-1*^*cKO*^ mice were significantly reduced (Fig. 5D). The loss of Bergmann glia during the period of CGN migration account for the observed cerebellar migration defects. Co-deletion of either one copy of *Bak* in *Mcl-1*^*cKO*^*/Bak*^*+/-*^ mice or both copies of *Bax* in *Mcl-1/Bax*^*dKO*^ mice was sufficient to rescue cerebellar migration, indicating that MCL-1 functions in Bergmann glia to restrict an apoptotic program that requires both BAX and BAK in order to drive cell death.

### *Mcl-1* deletion increases astrocytic populations

While *Mcl-1*^*cKO*^ mice contained fewer Bergmann glia, we noted increased numbers of GFAP+ astrocytic processes throughout the brain, most conspicuously in the relatively hypocellular molecular layer of the cerebellum (Fig. 6A). To determine if this increase in GFAP+ cells indicated an increase in the total number of astrocytes or an increase in the fraction of astrocytes expressing GFAP, we compared the number of cells expressing the astrocytic nuclear marker SOX9 in *Mcl-1*^*cKO*^ mice and controls (Fig. 6B,C). SOX9+ cells were not significantly increased in *Mcl-1*^*cKO*^ mice in any brain region at either P7 or P14, and rather were decreased in the corpus callosum at P14 (Fig. 6B,C). The increase in GFAP+ processes in *Mcl-1*^*cKO*^ mice without the increase in SOX9+ cells shows the *Mcl-1* deletion did not increase the total astrocyte population, but rather increased the fraction of astrocytes expressing GFAP, indicating a reactive response to injury (25, 26).

**Figure 6.**
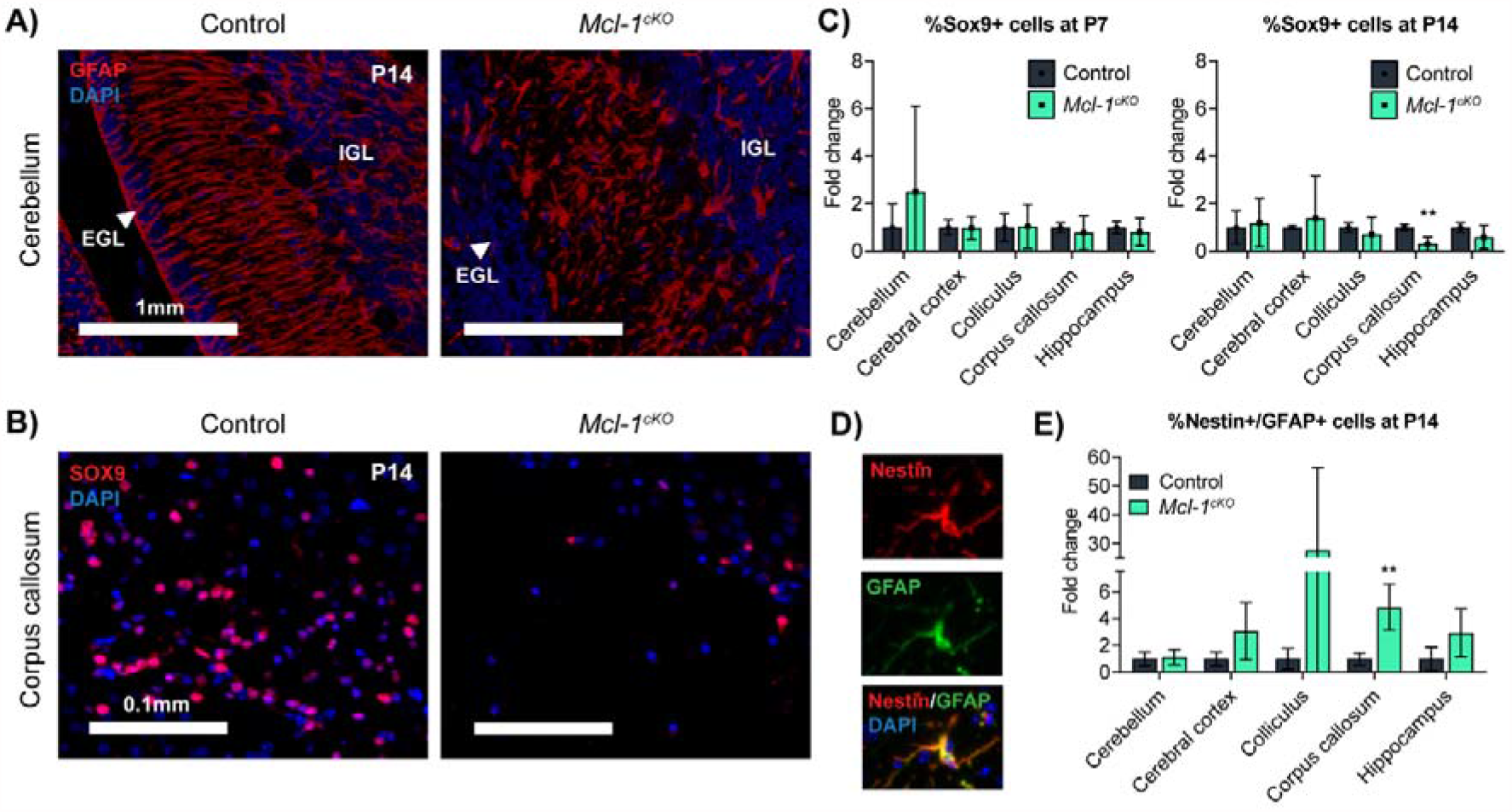
*Mcl-1* deletion causes reactive astrocytosis. (**A**) Representative GFAP IHC in sections of P14 cerebella in the indicated genotypes shows Bergmann glia processes in the control mice that are absent in the *Mcl-1*^*cKO*^, and numerous astrocyte processes in the *Mcl-1*^*cKO*^ mice. (**B**) Representative SOX9 IHC in sections of P14 corpus callosum in the indicated genotypes show a significant decrease of SOX9+ cells in the *Mcl-1*^*cKO*^ mice. (**C**) Quantification of SOX9 IHC at the indicated ages and genotypes, showing that astrocyte numbers are not increased the following scoring systemin *Mcl-1*^*cKO*^ mice. (**D**,**E**) Nestin/GFAP IHC, with representative images and quantification in the indicated genotypes. * and ** denote p<0.05 and p<0.01 respectively, relative to controls.

The intermediate filament protein NESTIN is known to be up-regulated in reactive astrocytes (27-30), and prior studies report increased Nestin-expressing astrocytes in white matter from humans with VWMD (31), and in astrocytes of the *eIF2B5* ^*R191H/R191H*^ mouse model of VWMD (8, 32). To determine if astrocytes in *Mcl1*-deleted mice had increased NESTIN expression, as seen in these leukodystrophies, we compared Nestin+/GFAP+ populations in P14 *Mcl-1*^*cKO*^ mice and controls (Fig. 6D). *Mcl-1*^*cKO*^ mice demonstrated increased fractions of Nestin+/GFAP+ cells in the corpus callosum (Fig. 6E), with a trend toward increased fractions in other white matter regions. *Mcl-1*^*cKO*^ mice showed astrocytic and oligodendrocytic abnormalities that are consistent with clinical white matter disease in patients and mouse models.

### Mcl-1 deletion causes neuroinflammation

Brains of *Mcl-1*^*cKO*^ mice contained increased cells expressing the myeloid marker IBA1 (Fig. 7A), indicating microglial activation (33, 34). The microglial changes in *Mcl-1*^*cKO*^ mice began later than the astrocytic and oligodendrocytic changes, beginning at the corpus callosum at P14; by P21, *Mcl-1*^*cKO*^ mice showed increased IBA+ cells in diverse brain regions (Fig. 7B). The increased IBA1+ microglia in *Mcl-1*^*cKO*^ mice demonstrates a non-cell autonomous effect in which *Mcl-1* deletion in the glio-neuronal GFAP lineage induced inflammatory changes in myeloid cells. This microglial activation is not seen in autopsy studies of VWMD patients, which are typically performed late in the disease process (35), but are seen in the VWMD mouse model, where brain samples can be studied earlier in the pathogenic process (36).

**Figure 7.**
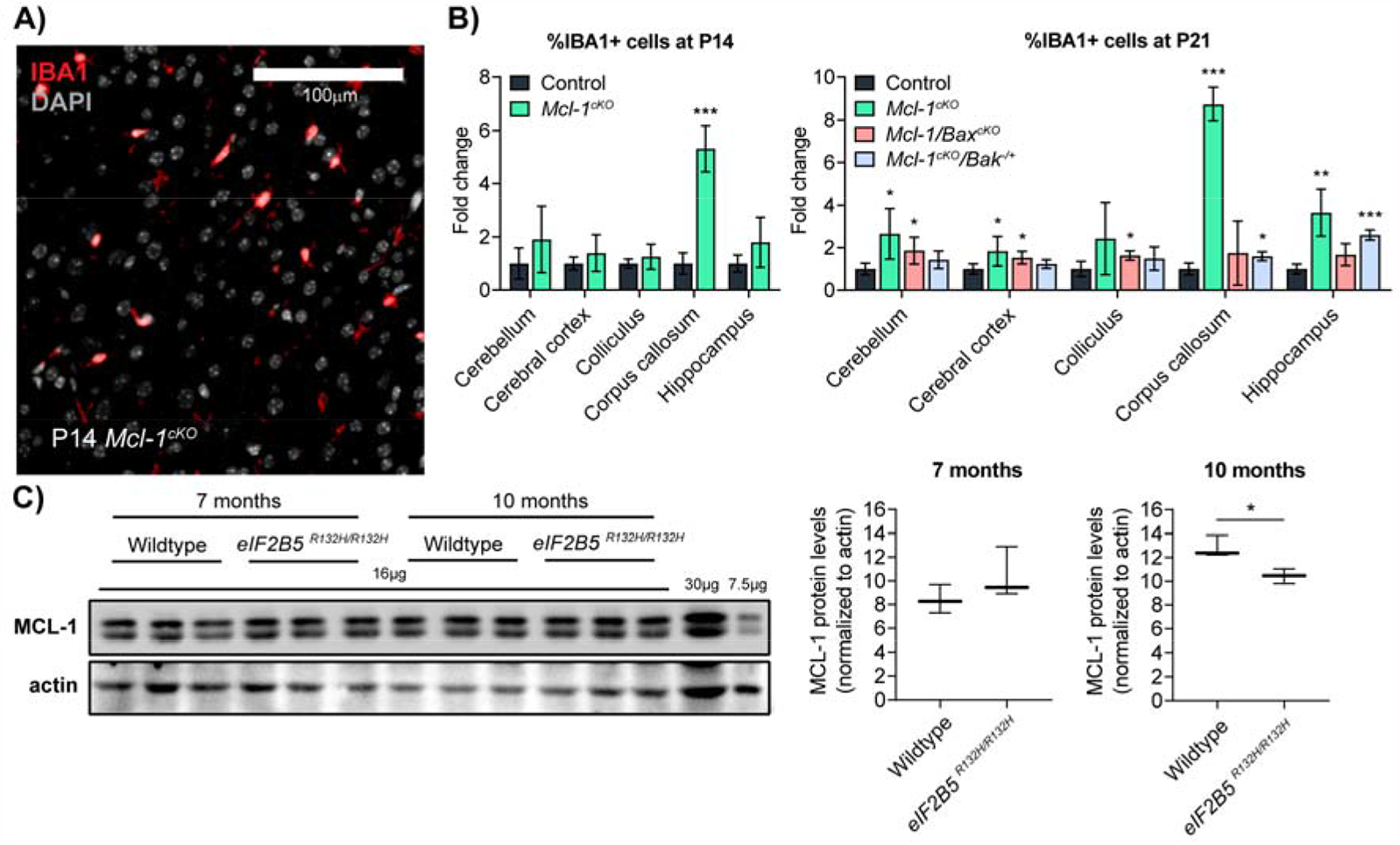
Progressive neuroinflammation in *Mcl-1*^*cKO*^ mice and reduced MCL-1 protein in mice with VWMD. (**A**,**B**) IBA1 IHC, with representative images and quantification in the indicated genotypes, showing a progressive increase in microglia in *Mcl-1*^*cKO*^ mice, beginning in the corpus callosum at P14 and expanding to include diverse regions by P21. Co-deletion of *Bak* or *Bax* variably rescue microgliosis with different effects in different regions. *, ** and *** denote p<0.05, p<0.01 and p<0.001 respectively, relative to controls. (**C**) Western blot comparing MCL-1 protein abundance in brain lysates of 7- and 10-months old WT and *eIF2B5*^*R132H/R132H*^ mice, demonstrating reduced MCL-1 in 10-month old mutant mice. * denotes p<0.03, relative to controls.

Co-deletion of either *Bak* or *Bax* prevented microgliosis in different brain regions. Heterozygous co-deletion of *Bak* in *Mcl-1*^*cKO*^*/Bak*^*+/-*^ mice significantly reduced IBA1+ cells in the cerebellum, cerebral cortex, colliculus, and corpus callosum, relative to *Mcl-1*^*cKO*^ mice. In contrast, homozygous co-deletion of *Bax* in *Mcl-1*^*cKO*^*/Bax*^*dKO*^ mice showed a significant decrease in IBA1+ cells in the hippocampus, relative to both *Mcl-1*^*cKO*^ and *Mcl-1*^*cKO*^*/Bak*^*+/-*^ mice (Fig. 7B). The rescue of neuro-inflammation by co-deletion of *Bax* or *Bak* implicates increased regional apoptosis as the cause of inflammation, while the region-specific pattern of rescue further supports the conclusion that white matter cells in different areas of the brain alternatively regulate apoptosis through interactions of MCL-1 with either BAX or BAK.

### Decreased brain MCL-1 protein in a VWMD mouse model

Based on the similarity of the *Mcl-1*^*cKO*^ phenotype to human leukodystrophies, we investigated whether MCL-1 expression is altered in the *eIF2B5* ^*R132H/R132H*^ mouse model of VWMD. We focused on this leukodystrophy model because MCL-1 abundance is known to depend on translational regulation and activation of the integrated stress response (ISR) (37), and both translation and ISR activation are abnormal in *eIF2B5* ^*R132H/R132H*^ mice (36, 38, 39). We compared MCL-1 abundance in brain lysates from wildtype and *eIF2B5* ^*R132H/R132H*^ mice at the pre-symptomatic time point of 7 months old and at 10 months old, when these mice begin to show phenotypic changes. At 7 months, MCL-1 abundance was similar, but at 10 months *eIF2B5* ^*R132H/R132H*^ mice showed reduced MCL-1 protein (Fig. 7C). The temporal correlation of reduced MCL-1 protein with the onset of brain pathology, suggests that MCL-1-regulated apoptosis may contribute to white matter pathology in *eIF2B5* ^*R132H/R132H*^ mice. Studies of Bak deletion in *eIF2B5* ^*R132H/R132H*^ mice will be needed to test directly for a role of apoptosis in VWMD pathology, as these data suggest.

## Discussion

Our data show that conditional deletion of anti-apoptotic *Mcl-1* in neuro-glial stem cells results in a progressive encephalopathy with post-natal white matter degeneration. While prior studies have shown increased apoptosis in embryonic and adult neural progenitors resulting from *Mcl-1* deletion (3, 40, 41), our data show that mice with brain-wide *Mcl-1* deletion are viable at birth and show normal brain anatomy through P7. In the postnatal brain of *Mcl-1*^*cKO*^ mice, however, differentiating oligodendrocytes underwent spontaneous apoptosis, resulting in impaired myelination and white matter degeneration, with preserved astrocyte populations. Bergmann glia also degenerated, disrupting granule neuron migration; the presence of both appropriately guided granule neurons in the IGL and ectopic granule neurons confirms that Bergmann glia were present in the early postnatal period and lost during the granule neuron migration period. While astrocyte populations were not decreased, astrocytes showed altered phenotypes, with increased expression of NESTIN and GFAP. Microglia were also increased and the microglial and astrocytic changes together suggest a reactive process, set in motion by apoptosis of MCL-1-dependent cell types.

The rescue of the *Mcl-1*^*cKO*^ phenotype by co-deletion of *Bak* or *Bax* implicate apoptosis as the process causing white matter loss in *Mcl-1*^*cKO*^ mice. These studies show that oligodendrocytes are primed for apoptosis and depend on MCL-1 protein to prevent inappropriate triggering of cell death programs. This apoptotic priming may contribute to the pathogenesis of diverse disorders that disproportionately affect CNS white matter.

Our data show that MCL-1 dependence is not homogeneous across glial subtypes, as *Mcl-1*-deleted astrocyte and OPC populations remain stable in size. Moreover, the different rescue effects seen with co-deletion of *Bax* or *Bak* in *Mcl-1*-deleted mice demonstrate that myelination defects and white matter loss can be dissociated. Co-deletion of *Bak* rescued both myelination, mature oligodendrocyte populations, and white matter loss. In contrast co-deletion of *Bax* reduced white matter loss without rescuing myelination or mature oligodendrocyte populations. These differences show that different types of glial cells with different, MCL-1-dependent mechanisms of apoptosis regulation, contribute to white matter development and stability and to the pathology of the *Mcl-1*^*cKO*^ phenotype.

Maintaining different thresholds for BAK-dependent and BAX-dependent apoptosis does not require different patterns of *Bak* and *Bax* expression. Prior studies show that both BAX and BAK drive apoptosis in cerebellar granule neuron progenitors, under different conditions. These cells undergo BAX-dependent apoptosis after radiation-induced DNA damage (42), but BAK-dependent apoptosis requires the more intense trigger of *Atr* deletion and resulting chromosomal fragmentation (11). Thus brain cells that express both BAX and BAK can show different sensitivities for triggering BAX-dependent or BAK-dependent apoptosis. Similarly, specific subsets of glia may regulate BAX-dependent and BAK-dependent apoptosis through different mechanisms. Our BAK rescue studies show that myelinating oligodendrocytes regulate apoptosis specifically through MCL-1:BAK interactions. The more limited rescue pattern with *Bax* co-deletion, which prevented white matter degeneration without normalizing myelination, demonstrates that a different type of glial cell requires both BAK and BAX to be present for *Mcl-1* deletion to produce phenotype changes. Thus, different types of glial cells depend on MCL-1 to regulate different pro-apoptotic proteins, and the loss of this regulation reproduces different aspects of the overall pathology. The loss of multiple subsets of glial cells may similarly contribute to different aspects of the leukodystrophy pathology.

Oligodendrocyte-specific apoptosis is observed in diverse leukodystrophies, including Pelizaeus-Merzbacher Disease (43), Pelizaeus-Merzbacher-like Disease (44), VWMD (45), and in the *twitcher* mouse model of globoid cell leukodystrophy (46). In each of these disorders, different mutations trigger different pathogenic processes with a final common result of inducing apoptosis in the oligodendrocyte lineage. Moreover, in these diverse disorders, apoptosis occurs as oligodendrocyte precursors differentiate into myelinating oligodendrocytes, as we found in *Mcl-1*^*cKO*^ mice. We propose that MCL-1 dependence contributes to the vulnerability of differentiating oligodendrocytes in diverse white matter disease states. In this context, our rescue studies show that blocking BAK or BAX can enhance oligodendrocyte survival, suggesting that treatments that inhibit the intrinsic apoptotic pathway in the postnatal brain may provide an avenue for reducing the impact of diverse leukodystrophies. Recent studies in the jimpy mouse model of Pelizaeus-Merzbacher disease support the potential for blocking apoptosis to block disease progression, as blocking ferroptosis through iron chelation improved myelination (43). Similarly, intracerebral treatment of jimpy mice with antisense oligonucleotides that inhibit PLP1 expression rescued white matter degeneration (47) and blocking BAK and BAX expression with similar methods may treat diverse white matter-specific disorders.

## Acknowledgments

supported by R01NS088219 (TRG), R01NS102627 (TRG), and R01NS1062 (TRG), T32CA071341 (AHC), R35GM128915 (VG), and R21CA227483 (VG), U54HD079124 (VDM,SM).

## Author Contributions

Conceptualization, T.R.G. and V.G.; Methodology, A.H.C., V.D.M., S.M, T.R.G., V.G. and O.E-S.; Investigation, A.H.C, V.D.M., A.R., L.A.A., M.H.,; Writing – Original Draft, A.H.C. and T.R.G.; Writing – Review & Editing, O.E-S., S.M., V.G. and T.R.G; Funding Acquisition, V.G., and T.R.G.;; Resources, O.E-S., V.G., and T.R.G.; Supervision, S.M.,O.E-S., V.G., and T.R.G.

## Conflicts of Interest

the authors have no competing financial interests in relation to the work described.

## Supplementary Figures

**Supplementary Figure 1.**
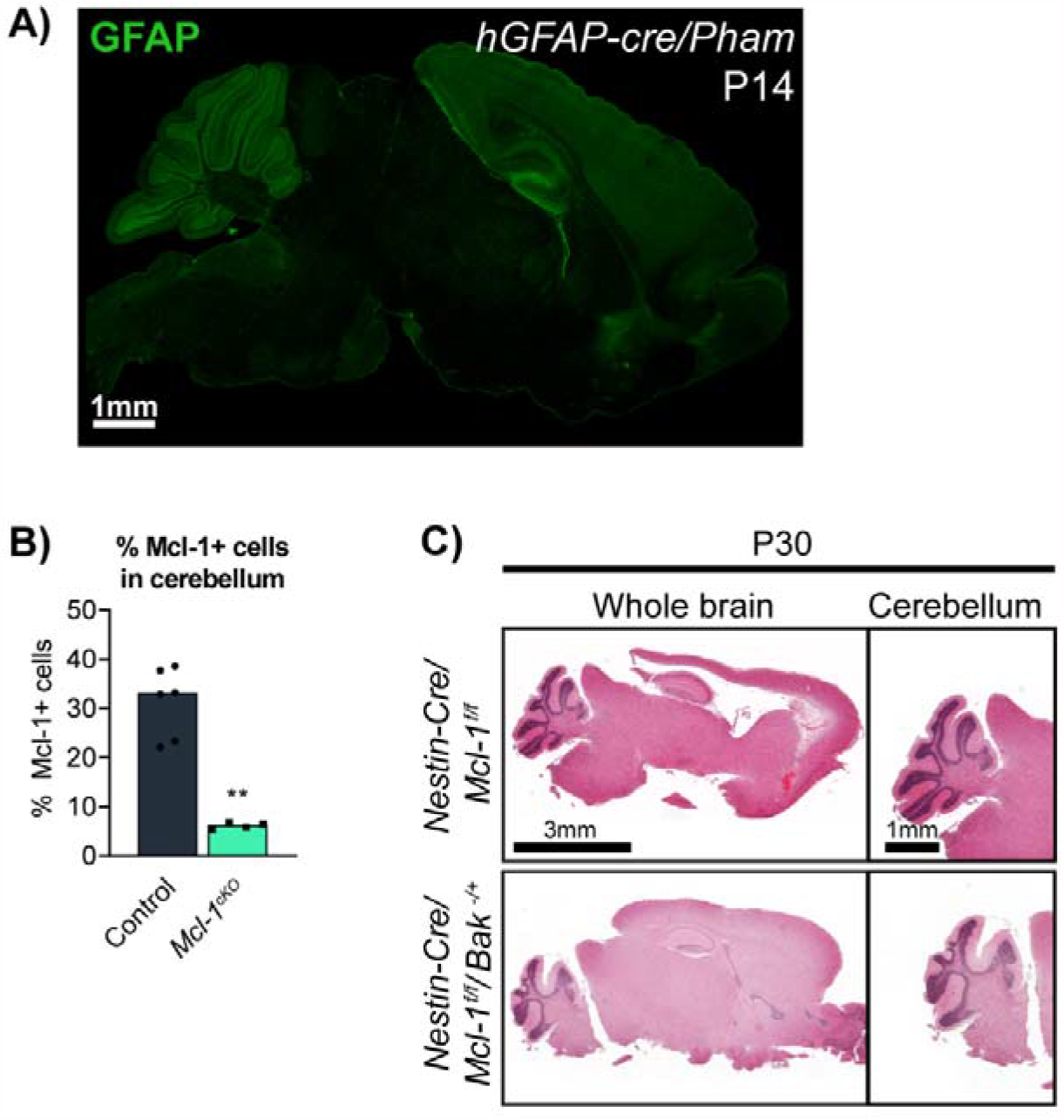
*hGFAP-Cre* conditionally deletes *Mcl-1* in regions throughout the brain and conditional Mcl-1 deletion driven by *Nestin-Cre* produces a similar phenotype. **A)** *hGFAP-Cre* induces expression of the fluorescent reporter DENDRA2 in *hGFAP-cre/Pham* mice, indicating regions of GFAP expression. *hGFAP-Cre* targets cells in diverse regions throughout the brain, including the cerebellum, hippocampus, and cortex. **B)** MCL-1 protein levels are significantly reduced in cerebella of *Mcl-1*^*cKO*^ mice. **C)** Like *Mcl-1*^*cKO*^ mice, *Nestin-Cre/Mcl-1*^*loxP/loxP*^ mice showed white matter degeneration that was rescued by heterozygous *Bak* co-deletion.

**Supplementary Figure 2.**
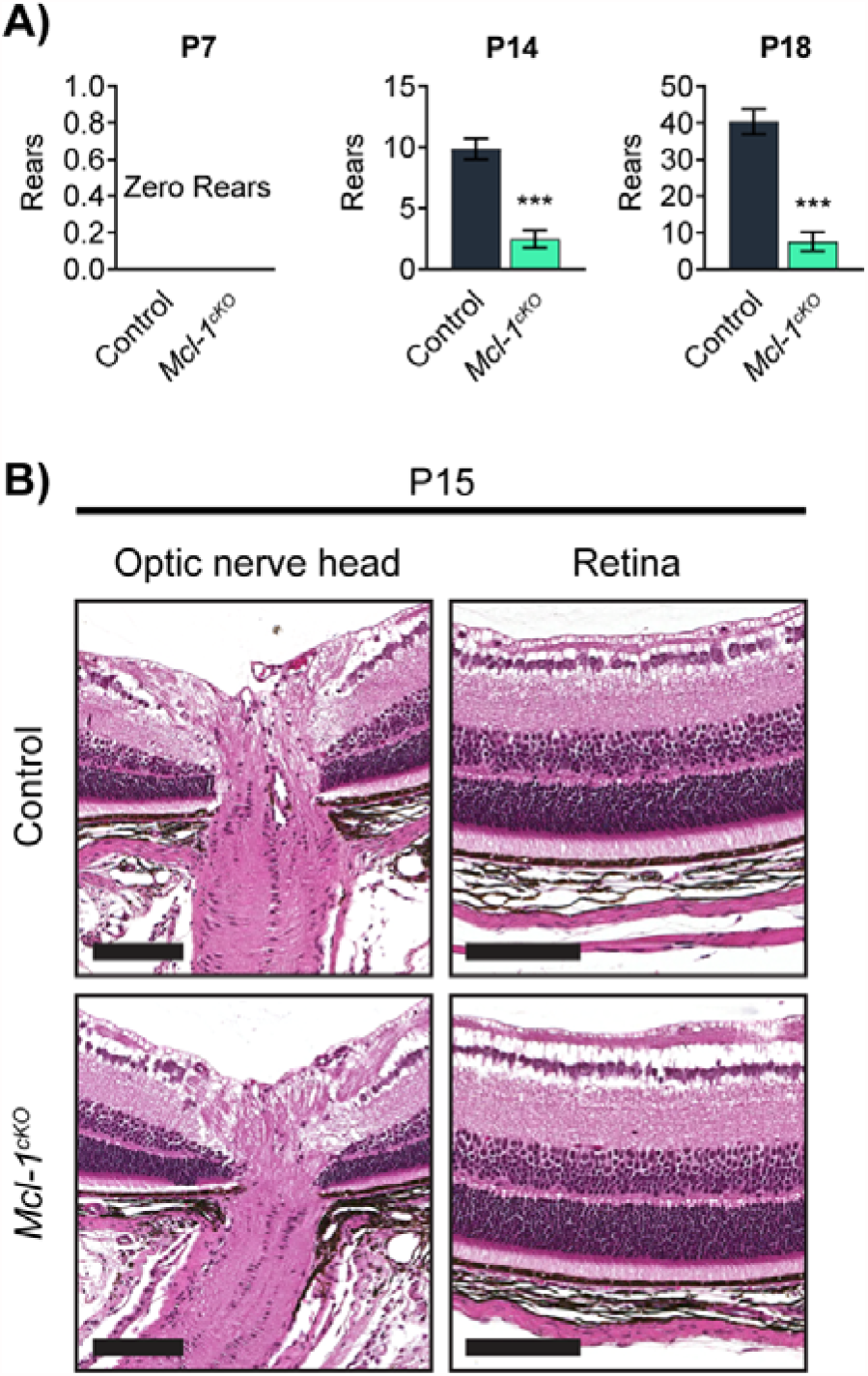
Open field rearing behavior changes and absence of retinal pathology in *Mcl-1*^*cKO*^ mice. **A)** Decreases rearing in *Mcl-1*^*cKO*^ mice at P14 and P21. **B)** Normal-appearing retina in *Mcl-1*^*cKO*^ mice at P15, when brain pathology is fulminant. Scale bar = 0.1mm.

**Supplementary Figure 3.**
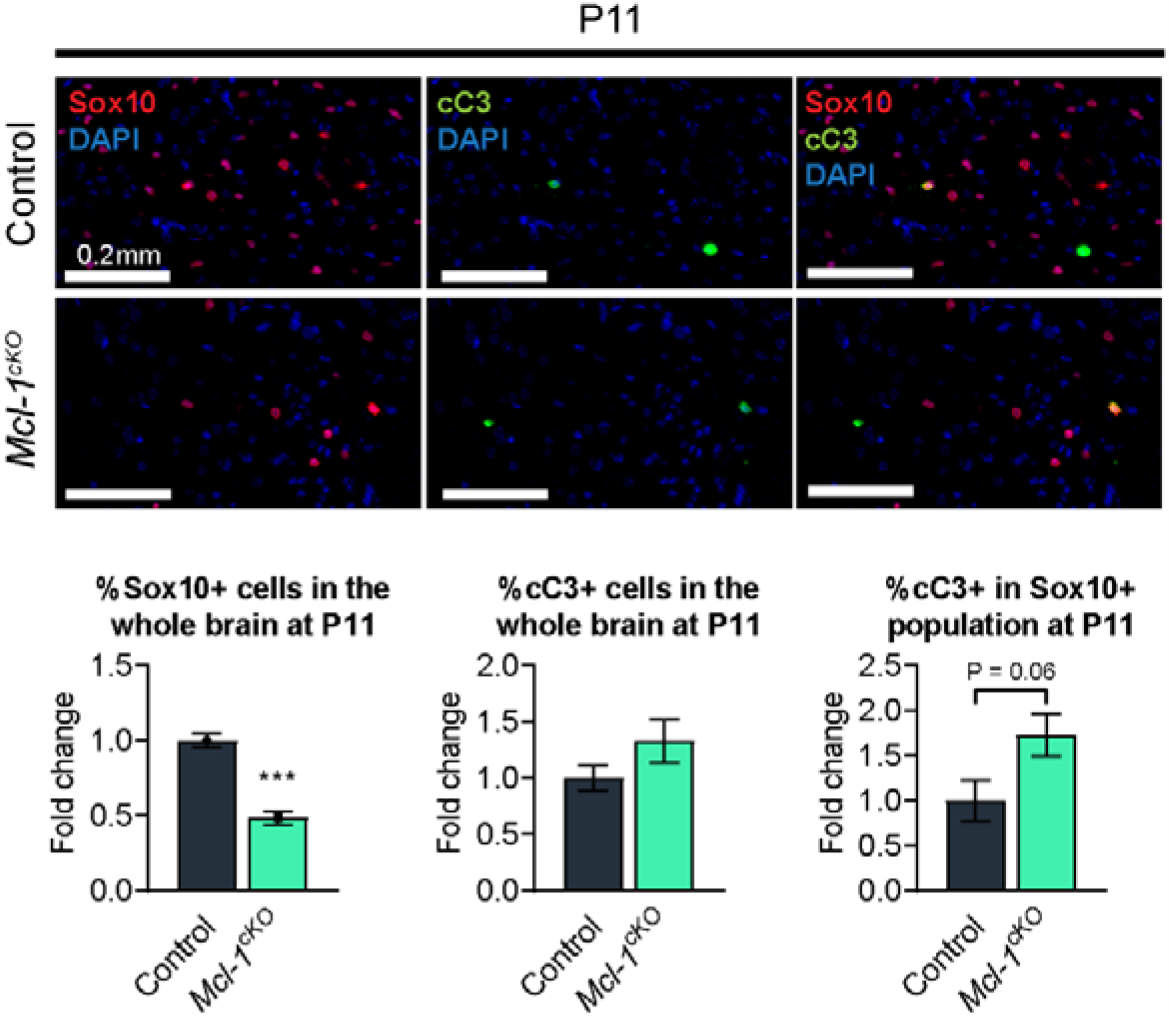
Immunofluorescence for cC3 and SOX10 in *Mcl-1*^*cKO*^ mice and controls at P11. SOX10 is significantly decreased, but no statistically significant changes are noted in the total cC3+ cells or cC3+ cells within the SOX10+ subset.

## Notes

### Competing Interest Statement

The authors have declared no competing interest.

### Summary of Updates

This revision includes a complete re-writing of the text to focus on apoptosis regulation. A potential role for MCL-1 in Vanishing White Matter disease has been de-emphasized.

## References

1. Hollville E, Romero SE, Deshmukh M. Apoptotic cell death regulation in neurons. FEBS J. 2019;286(17):3276–98.

2. Kole AJ, Annis RP, Deshmukh M. Mature neurons: equipped for survival. Cell Death Dis. 2013;4:e689.

3. Fogarty LC, Flemmer RT, Geizer BA, Licursi M, Karunanithy A, Opferman JT, et al. Mcl-1 and Bcl-xL are essential for survival of the developing nervous system. Cell Death Differ. 2019;26(8):1501–15.

4. Nakamura A, Swahari V, Plestant C, Smith I, McCoy E, Smith S, et al. Bcl-xL Is Essential for the Survival and Function of Differentiated Neurons in the Cortex That Control Complex Behaviors. J Neurosci. 2016;36(20):5448–61.

5. Veleta KA, Cleveland AH, Babcock BR, He YW, Hwang D, Sokolsky-Papkov M, et al. Antiapoptotic Bcl-2 family proteins BCL-xL and MCL-1 integrate neural progenitor survival and proliferation during postnatal cerebellar neurogenesis. Cell Death Differ. 2020.

6. Crowther AJ, Gama V, Bevilacqua A, Chang SX, Yuan H, Deshmukh M, et al. Tonic activation of Bax primes neural progenitors for rapid apoptosis through a mechanism preserved in medulloblastoma. The Journal of neuroscience : the official journal of the Society for Neuroscience. 2013;33(46):18098–108.

7. Dzhagalov I, St John A, He YW. The antiapoptotic protein Mcl-1 is essential for the survival of neutrophils but not macrophages. Blood. 2007;109(4):1620–6.

8. Geva M, Cabilly Y, Assaf Y, Mindroul N, Marom L, Raini G, et al. A mouse model for eukaryotic translation initiation factor 2B-leucodystrophy reveals abnormal development of brain white matter. Brain. 2010;133(Pt 8):2448–61.

9. Ocasio JK, Bates RDP, Rapp CD, Gershon TR. GSK-3 modulates SHH-driven proliferation in postnatal cerebellar neurogenesis and medulloblastoma. Development. 2019;146(20).

10. Tech K, Tikunov AP, Farooq H, Morrissy AS, Meidinger J, Fish T, et al. Pyruvate Kinase Inhibits Proliferation during Postnatal Cerebellar Neurogenesis and Suppresses Medulloblastoma Formation. Cancer Res. 2017;77(12):3217–30.

11. Lang PY, Nanjangud GJ, Sokolsky-Papkov M, Shaw C, Hwang D, Parker JS, et al. ATR maintains chromosomal integrity during postnatal cerebellar neurogenesis and is required for medulloblastoma formation. Development. 2016;143(21):4038–52.

12. Crowther AJ, Ocasio JK, Fang F, Meidinger J, Wu J, Deal AM, et al. Radiation Sensitivity in a Preclinical Mouse Model of Medulloblastoma Relies on the Function of the Intrinsic Apoptotic Pathway. Cancer Res. 2016;76(11):3211–23.

13. Borras T, Cowley DO, Asokan P, Pandya K. Generation of a Matrix Gla (Mgp) floxed mouse, followed by conditional knockout, uncovers a new Mgp function in the eye. Sci Rep. 2020;10(1):18583.

14. Williams SE, Garcia I, Crowther AJ, Li S, Stewart A, Liu H, et al. Aspm sustains postnatal cerebellar neurogenesis and medulloblastoma growth. Development. 2015.

15. Cheh MA, Millonig JH, Roselli LM, Ming X, Jacobsen E, Kamdar S, et al. En2 knockout mice display neurobehavioral and neurochemical alterations relevant to autism spectrum disorder. Brain Res. 2006;1116(1):166–76.

16. Zhuo L, Theis M, Alvarez-Maya I, Brenner M, Willecke K, Messing A. hGFAP-cre transgenic mice for manipulation of glial and neuronal function in vivo. Genesis. 2001;31(2):85–94.

17. Kuang Y, Liu Q, Shu X, Zhang C, Huang N, Li J, et al. Dicer1 and MiR-9 are required for proper Notch1 signaling and the Bergmann glial phenotype in the developing mouse cerebellum. Glia. 2012;60(11):1734–46.

18. Guyenet SJ, Furrer SA, Damian VM, Baughan TD, La Spada AR, Garden GA. A simple composite phenotype scoring system for evaluating mouse models of cerebellar ataxia. J Vis Exp. 2010(39).

19. Vanderver A, Tonduti D, Schiffmann R, Schmidt J, van der Knaap MS. Leukodystrophy Overview - ARCHIVED CHAPTER, FOR HISTORICAL REFERENCE ONLY. In: Adam MP, Ardinger HH, Pagon RA, Wallace SE, Bean Ljh, Stephens K, et al., editors. GeneReviews((R)). Seattle (WA)1993.

20. van der Knaap MS, Fogli A, Boespflug-Tanguy O, Abbink TEM, Schiffmann R. Childhood Ataxia with Central Nervous System Hypomyelination / Vanishing White Matter. In: Adam MP, Ardinger HH, Pagon RA, Wallace SE, Bean Ljh, Stephens K, et al., editors. GeneReviews((R)). Seattle (WA)1993.

21. Glasgow SM, Zhu W, Stolt CC, Huang TW, Chen F, LoTurco JJ, et al. Mutual antagonism between Sox10 and NFIA regulates diversification of glial lineages and glioma subtypes. Nat Neurosci. 2014;17(10):1322–9.

22. Nishiyama A, Boshans L, Goncalves CM, Wegrzyn J, Patel KD. Lineage, fate, and fate potential of NG2-glia. Brain Res. 2016;1638(Pt B):116–28.

23. Hatten ME. Riding the glial monorail: a common mechanism for glial-guided neuronal migration in different regions of the developing mammalian brain. Trends Neurosci. 1990;13(5):179–84.

24. Rakic P. Neuron-glia relationship during granule cell migration in developing cerebellar cortex. A Golgi and electronmicroscopic study in Macacus Rhesus. J Comp Neurol. 1971;141(3):283–312.

25. Eddleston M, Mucke L. Molecular profile of reactive astrocytes--implications for their role in neurologic disease. Neuroscience. 1993;54(1):15–36.

26. Dahl D, Bignami A. Heterogeneity of the glial fibrillary acidic protein in gliosed human brains. J Neurol Sci. 1974;23(4):551–63.

27. Tamagno I, Schiffer D. Nestin expression in reactive astrocytes of human pathology. J Neurooncol. 2006;80(3):227–33.

28. Schmidt-Kastner R, Aguirre-Chen C, Saul I, Yick L, Hamasaki D, Busto R, et al. Astrocytes react to oligemia in the forebrain induced by chronic bilateral common carotid artery occlusion in rats. Brain Res. 2005;1052(1):28–39.

29. Sahin Kaya S, Mahmood A, Li Y, Yavuz E, Chopp M. Expression of nestin after traumatic brain injury in rat brain. Brain Res. 1999;840(1-2):153–7.

30. Clarke SR, Shetty AK, Bradley JL, Turner DA. Reactive astrocytes express the embryonic intermediate neurofilament nestin. Neuroreport. 1994;5(15):1885–8.

31. Bugiani M, Boor I, van Kollenburg B, Postma N, Polder E, van Berkel C, et al. Defective glial maturation in vanishing white matter disease. J Neuropathol Exp Neurol. 2011;70(1):69–82.

32. Dooves S, Bugiani M, Postma NL, Polder E, Land N, Horan ST, et al. Astrocytes are central in the pathomechanisms of vanishing white matter. J Clin Invest. 2016;126(4):1512–24.

33. Ito D, Imai Y, Ohsawa K, Nakajima K, Fukuuchi Y, Kohsaka S. Microglia-specific localisation of a novel calcium binding protein, Iba1. Brain Res Mol Brain Res. 1998;57(1):1–9.

34. Imai Y, Ibata I, Ito D, Ohsawa K, Kohsaka S. A novel gene iba1 in the major histocompatibility complex class III region encoding an EF hand protein expressed in a monocytic lineage. Biochem Biophys Res Commun. 1996;224(3):855–62.

35. Rodriguez D, Gelot A, della Gaspera B, Robain O, Ponsot G, Sarlieve LL, et al. Increased density of oligodendrocytes in childhood ataxia with diffuse central hypomyelination (CACH) syndrome: neuropathological and biochemical study of two cases. Acta Neuropathol. 1999;97(5):469–80.

36. Wong YL, LeBon L, Basso AM, Kohlhaas KL, Nikkel AL, Robb HM, et al. eIF2B activator prevents neurological defects caused by a chronic integrated stress response. Elife. 2019;8.

37. Fritsch RM, Schneider G, Saur D, Scheibel M, Schmid RM. Translational repression of MCL-1 couples stress-induced eIF2 alpha phosphorylation to mitochondrial apoptosis initiation. J Biol Chem. 2007;282(31):22551–62.

38. Abbink TEM, Wisse LE, Jaku E, Thiecke MJ, Voltolini-Gonzalez D, Fritsen H, et al. Vanishing white matter: deregulated integrated stress response as therapy target. Ann Clin Transl Neurol. 2019;6(8):1407–22.

39. Moon SL, Parker R. EIF2B2 mutations in vanishing white matter disease hypersuppress translation and delay recovery during the integrated stress response. RNA. 2018;24(6):841–52.

40. Hasan SM, Sheen AD, Power AM, Langevin LM, Xiong J, Furlong M, et al. Mcl1 regulates the terminal mitosis of neural precursor cells in the mammalian brain through p27Kip1. Development. 2013;140(15):3118–27.

41. Malone CD, Hasan SM, Roome RB, Xiong J, Furlong M, Opferman JT, et al. Mcl-1 regulates the survival of adult neural precursor cells. Mol Cell Neurosci. 2012;49(4):439–47.

42. Chong MJ, Murray MR, Gosink EC, Russell HRC, Srinivasan A, Kapsetaki M, et al. Atm and Bax cooperate in ionizing radiation-induced apoptosis in the central nervous system. Proceedings of the National Academy of Sciences. 2000;97(2):889–94.

43. Nobuta H, Yang N, Ng YH, Marro SG, Sabeur K, Chavali M, et al. Oligodendrocyte Death in Pelizaeus-Merzbacher Disease Is Rescued by Iron Chelation. Cell Stem Cell. 2019;25(4):531–41 e6.

44. Georgiou E, Sidiropoulou K, Richter J, Papaneophytou C, Sargiannidou I, Kagiava A, et al. Gene therapy targeting oligodendrocytes provides therapeutic benefit in a leukodystrophy model. Brain. 2017;140(3):599–616.

45. Van Haren K, van der Voorn JP, Peterson DR, van der Knaap MS, Powers JM. The life and death of oligodendrocytes in vanishing white matter disease. J Neuropathol Exp Neurol. 2004;63(6):618–30.

46. Taniike M, Mohri I, Eguchi N, Irikura D, Urade Y, Okada S, et al. An apoptotic depletion of oligodendrocytes in the twitcher, a murine model of globoid cell leukodystrophy. J Neuropathol Exp Neurol. 1999;58(6):644–53.

47. Elitt MS, Barbar L, Shick HE, Powers BE, Maeno-Hikichi Y, Madhavan M, et al. Suppression of proteolipid protein rescues Pelizaeus-Merzbacher disease. Nature. 2020;585(7825):397–403.

